# Context-specific effects of facial dominance and trustworthiness on hypothetical leadership decisions

**DOI:** 10.1101/575597

**Authors:** Hannah S. Ferguson, Anya Owen, Amanda C. Hahn, Jaimie Torrance, Lisa M. DeBruine, Benedict C. Jones

## Abstract

Social judgments of faces predict important social outcomes, including leadership decisions. Previous work suggests that facial cues associated with perceptions of dominance and trustworthiness have context-specific effects on leadership decisions. Facial cues linked to perceived dominance have been found to be preferred in leaders for hypothetical wartime contexts and facial cues linked to perceived trustworthiness have been found to be preferred in leaders for hypothetical peacetime contexts. Here we sought to replicate these effects using images of women’s faces. Consistent with previous work, a linear mixed effects model demonstrated that more trustworthy-looking faces were preferred in leaders during times of peace and more dominant-looking faces were preferred in leaders during times of war. These results provide converging evidence for context-specific effects of facial cues on hypothetical leadership judgments.

## Introduction

Social judgments of faces predict important social outcomes, such as romantic and platonic partner choices and hiring decisions (e.g., Rhodes, 2006; Todorov et al., 2015). One area that has received considerable attention in the social perception literature is the role that social judgments of faces play in hypothetical leadership decisions (reviewed in Van Vugt & Grabo, 2015). Indeed, several lines of evidence suggest that even very rapid social judgments of politicians’ faces predict actual election outcomes (Antonakis & Dalgas, 2009; Little et al., 2007; Todorov et al., 2005).

Several studies have found that people whose faces are judged to look particularly dominant or trustworthy are preferred in hypothetical leadership decisions (see Little et al., 2012). However, other research suggests that facial characteristics linked to perceptions of dominance and trustworthiness can have context-specific effects on hypothetical leadership decisions (Little et al., 2012; Van Vugt & Grabo, 2015). People with masculine faces are generally perceived to look more dominant (Perrett et al., 1998) and tend to be preferred as leaders in hypothetical wartime scenarios (Spisak et al., 2012). By contrast, people with feminine faces are generally perceived to look more trustworthy (Perrett et al., 1998) and tend to be preferred as leaders in hypothetical peacetime scenarios (Spisak et al., 2012). These results have been interpreted as evidence that stereotypic perceptions of candidates’ suitability for particular types of leadership roles influence hypothetical leadership decisions, potentially reflecting the context-specific relevance of these traits for different types of coalitions (Spisak et al., 2012; Van Vugt & Grabo, 2015; Little et al., 2007).

Spisak et al. (2012) demonstrated context-specific effects of facial appearance on leadership decisions using face stimuli that had been experimentally manipulated along a masculinity-femininity dimension. Some researchers have criticized this method (i.e., experimental manipulation of facial characteristics) because the observed effects on perceptions may not generalize well to judgments of natural face images that vary simultaneously on many dimensions (e.g., Scott et al., 2010). Because of such criticisms, we attempted to conceptually replicate Spisak et al’s results for context-specific leadership judgments using dominance and trustworthiness ratings of unmanipulated face images. Unmanipulated female faces, varying naturally in perceived dominance and trustworthiness, were judged for leadership ability in a hypothetical peacetime or wartime scenario. Given Spisak et al’s (2012) findings, we predicted that more dominant-looking faces would be preferred in leaders during the wartime context, while more trustworthy-looking faces would be preferred in leaders during the peacetime context.

## Methods

One hundred men (mean age=26.22 years, SD=6.11 years) and 100 women (mean age=24.71 years, SD=5.4 years) were randomly allocated to rate face images of 50 young adult white women for either trustworthiness or dominance using 1 (very low) to 7 (very high) scales. Inter-rater agreement was high for both sets of ratings (both Cronbach’s alphas > .90) A different group of 137 men (mean age=29.4 years, SD=10.91 years) and 237 women (mean age=25.45 years, SD=9.43 years) were randomly allocated to rate the same 50 female faces for leadership at a time of war or leadership at a time of peace, also using a 1 (very bad) to 7 (very good) scale. The specific questions asked for leadership ratings were “How good a leader would this person be for a country during a time of war?” and “How good a leader would this person be for a country during a time of peace?”. Trial order was fully randomized for all ratings.

## Results

Data were analyzed using a linear mixed effects model with leadership rating as the dependent variable and average dominance rating, average trust rating, and leadership context as predictors. The model also included interactions between dominance rating and leadership context and between trustworthiness rating and leadership context. Random intercepts were specified for the 50 faces and the 373 participants. Random slopes were specified maximally (dominance and trustworthiness ratings by participant and context by face). Data and analysis code are publicly available at https://osf.io/q54nm/.

The relationship between leadership ratings and dominance ratings was qualified by context (beta = 0.38, SE = 0.08, t(149.99) = 4.59, p < .001, 95% CI = [0.22, 0.55]). The relationship between leadership ratings and trustworthiness ratings was also qualified by context (beta = −0.4, SE = 0.08, t(129.37) = −4.93, p < .001, 95% CI = [−0.55, −0.24]). As predicted, dominance ratings were positively related to leadership ratings in the war, but not peace, context, while trustworthiness ratings were positively related to leadership ratings in the peace, but not war, context (see Figure 1).

**Figure 1.**
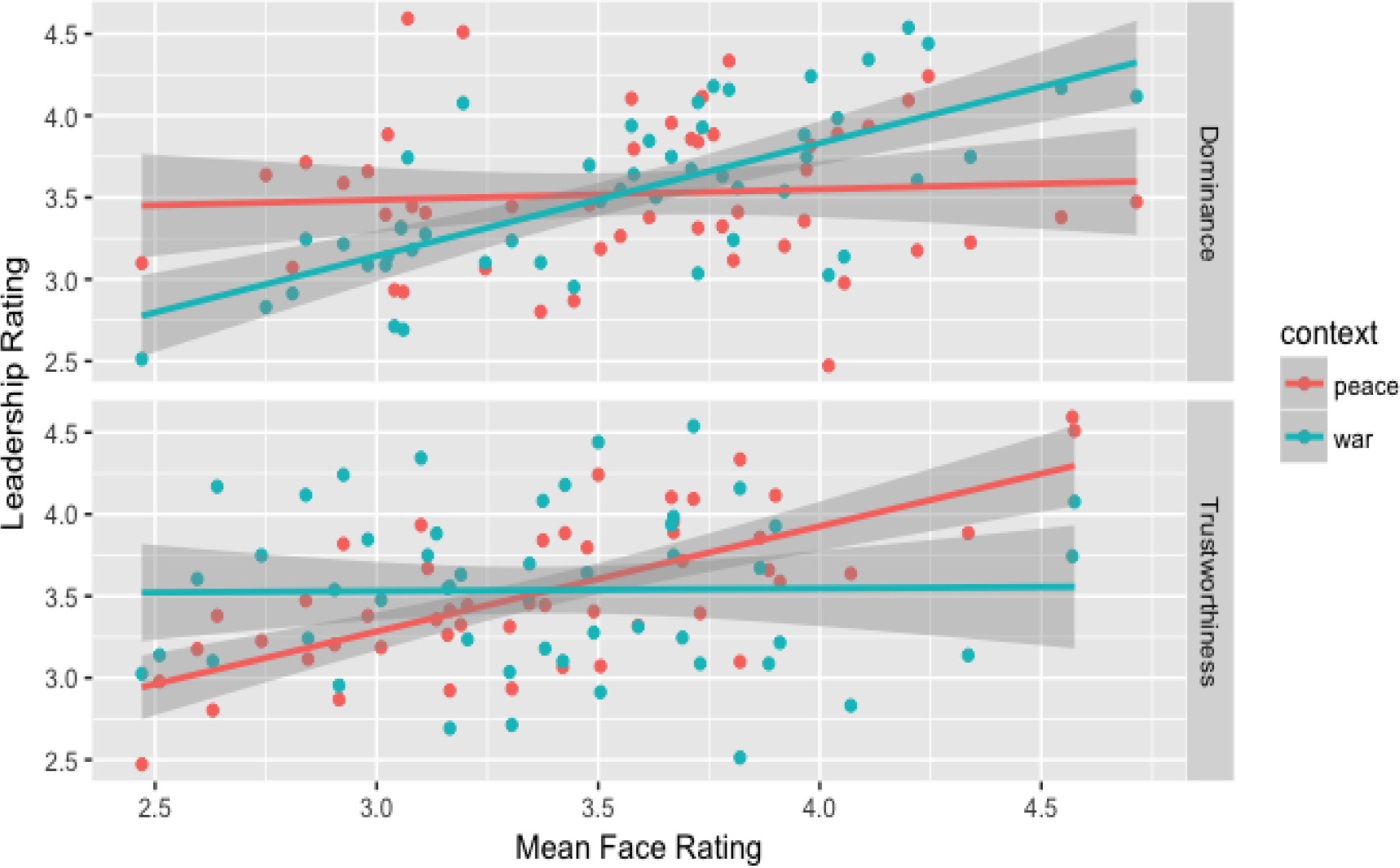
Context-specific effects of facial dominance and trustworthiness on leadership judgements.

## Discussion

The current study investigated hypothetical leadership decisions using female faces that varied naturally in perceived dominance and trustworthiness for two different leadership contexts; peacetime and wartime. Consistent with Spisak et al’s (2012) findings, our results indicate that women with more dominant-looking faces were preferred as leaders during hypothetical wartime contexts, while women with more trustworthy-looking faces were preferred as leaders during hypothetical peacetime contexts.

While previous research has focused primarily on leadership decisions of male faces, our results add to a growing body of research demonstrating that the effects of facial appearance on perceived leadership ability extend to female faces (Re et al. 2013; Van Vugt & Spisak, 2008). Although some previous research has suggested that manipulating the masculinity of female faces does not increase perceived dominance as much as it does in male face counterparts (Re et al, 2013; Watkins et al, 2010), several studies investigating the impact of facial cues on leadership decisions have demonstrated that cues of masculinity-femininity are more influential than actual sex cues in determining leadership decisions in hypothetical voting tasks (Carpinella, 2016; Spisak et al, 2012).

The pattern of results that we observed provides converging evidence for the hypothesis that facial cues have context-dependent effects on leadership decisions. Although evidence that perceptions of other peoples’ dominance and trustworthiness are accurate is mixed and controversial (Todorov et al., 2015), our findings present further evidence that stereotypic perceptions of faces shape social judgments in ways that are predictable and somewhat rational.

